# Assessing Brain-Behaviour Coupling in Non-invasive Brain Stimulation Using Reliable Change Indices: Evidence from pre-Supplementary Motor Area – right Inferior Frontal Gyrus transcranial Alternating Current Stimulation

**DOI:** 10.64898/2026.03.24.714072

**Authors:** Hakuei Fujiyama, Kym Wansbrough, Brooke Lebihan, Jane Tan, Oron Levin, Danielle C. Mathersul, Alexander D Tang

## Abstract

Non-invasive brain stimulation (NiBS) studies frequently report exploratory correlations between individual-level changes in neurophysiological and behavioural measures. However, these analyses are typically underpowered and rely on ratio-based change scores with known statistical limitations. We addressed these limitations by pooling individual data from three independent studies (total N = 69), providing adequate power to detect small-to-medium effects. All studies applied 20 Hz transcranial alternating current stimulation (tACS) targeting the pre-supplementary motor area (preSMA) and right inferior frontal gyrus (rIFG), regions central to inhibitory control. Changes in preSMA–rIFG connectivity measured with EEG imaginary coherence (ImCoh) and response inhibition (stop-signal reaction time, SSRT) were quantified using reliable change indices (RCI), which were z-standardised within studies to enable pooled mixed-effects regression. No meaningful association was found between tACS-induced ImCoh change and SSRT change (r = .013, marginal R² = .004), with project-wise correlations that were small, non-significant, and inconsistent in direction. Sensitivity analysis using ratio-based change scores converged on the same null result (r = .014), though ratio scores showed severe distributional violations relative to the approximately normal RCI distributions, supporting the methodological case for RCI on statistical grounds.

These results provide no support for a systematic individual-level brain–behaviour coupling between preSMA–rIFG connectivity and response inhibition following 20 Hz tACS, and suggest that any true effect is likely to be small. The present work offers a methodological benchmark for quantifying individual-level brain–behaviour coupling in NiBS research, and highlights the need for more sensitive neural markers and adequately powered design.

---

Non-invasive brain stimulation (NiBS) techniques such as transcranial magnetic stimulation (TMS) and transcranial electrical stimulation (tES) have emerged as promising tools for modulating neurophysiological activity and behaviour. However, the extent to which NiBS-induced neurophysiological modulations (e.g., motor-evoked potentials) reliably predict corresponding behavioural outcomes (e.g., improvements in reaction time) remains an open and critical question in the field. Establishing reliable brain behaviour relationships is key for identifying mechanistic markers of stimulation efficacy and for guiding the optimisation and translation of NiBS interventions (e.g., Bestmann & Krakauer, 2015; Ryan et al., 2023).

A number of studies reported NiBS-induced changes in cortical excitability (e.g., Fujiyama et al., 2014, 2017; Nitsche & Paulus, 2001), synaptic plasticity (e.g., Farahani et al., 2021; Tang et al., 2021; Vlachos et al., 2012), network connectivity (e.g., Lebihan et al., 2025; Reinhart & Nguyen, 2019; Wansbrough, Marinovic, et al., 2024), and cognitive/motor performance (e.g., Hummel et al., 2010; Lebihan et al., 2025; Zimerman et al., 2013). These findings show that NiBS can modulate neural and behavioural processes at the group level and have provided valuable insight into the causal role of the human cortex in behaviour (Ridding & Ziemann, 2010).

Moreover, these findings provide a justification to explore the potential therapeutic effects of NiBS in patients across diverse neurological and psychiatric disorders, including stroke, Parkinson’s disease, and major depressive disorder (for review, see Rektorová et al., 2025). However, establishing clear links between stimulation-induced neurophysiological changes and behavioural outcomes at the individual level has proven challenging. While the induction of lasting changes in cortical excitability can, under some conditions, modulate behaviour and interact with ongoing learning processes (e.g., Ridding & Ziemann, 2010), the extent to which measurable neurophysiological markers reliably predict behavioural change remains unclear. Clarifying this relationship is important for understanding stimulation mechanisms and optimising behavioural outcomes, and has motivated several studies to explore associations between neurophysiological and behavioural changes following NiBS. Several of these studies, which are included in a review conducted by Ryan and colleagues (2023), report exploratory post-hoc correlations between neurophysiological markers and behavioural outcomes, despite being primarily powered to detect group-level intervention effects. These exploratory analyses, while valuable for hypothesis generation, face inherent limitations: 1) sample sizes adequate for detecting main effects often lack statistical power for robust correlation analyses (Zehetmayer et al., 2015), 2) multiple unadjusted comparisons increase false discovery rates (Benjamini et al., 2001), and 3) readers may misinterpret nominally significant correlations as conclusive mechanistic evidence despite cautionary statements of the insufficient statistical power to detect the correlation. Detecting non-trivial correlations between neurophysiological and behavioural change likely requires substantially larger sample sizes than those typically used in studies aiming to detect pre-post intervention effects, highlighting the importance of adequate planning (Kiernan & Baiocchi, 2022).

Another limitation of brain-behaviour correlation analyses is that most studies used change scores (e.g., the ratio of pre- and post-assessment values) to account for baseline differences by standardising change. Although it is intuitively appealing, such ratio-based measures have undesirable statistical properties. This approach inadvertently increases sampling variation, such that ratio-based measures often exhibit substantially greater variability than the original variable, reducing statistical reliability (Jasieński & Bazzaz, 1999). Ratio values also frequently violate the assumptions of parametric tests, including normality and homoscedasticity, since the ratio of two normally distributed variables does not generally follow a normal distribution (Allison et al., 1995; Díaz-Francés & Rubio, 2013). Furthermore, the use of ratios can introduce spurious correlations, creating apparent associations between variables that are purely artefacts of the division process rather than reflecting genuine relationships (Kronmal, 1993). Despite these limitations, standardised ratios continue to be used in exploratory investigations of neural correlates in the NiBS field.

A statistically tractable alternative for quantifying change in repeated-measure designs is the reliable change index (RCI) (Jacobson & Truax, 1991). RCI standardises pre-post differences relative to the expected variability of the measure, thus accounting for measurement reliability. This yields a standardised change score for each individual, allowing change to be compared on a common scale and distinguishing true change from measurement error. RCI, therefore, provides a principled approach for evaluating whether stimulation-induced neurophysiological change is meaningfully associated with stimulation-induced behavioural change, while reducing the influence of noise-related variability that can attenuate correlations between neurophysiological and behavioural outcomes.

In the present study, we used RCIs as continuous indices of reliable change to conduct a coordinated analysis of archived datasets from our research group, combining individual data from three NiBS studies published between 2023 and 2025 (N = 69), and examining correlations between neurophysiological and behavioural RCIs. All three studies applied 20Hz transcranial alternating current stimulation (tACS), aiming to improve functional connectivity between the pre-supplementary motor area (preSMA) and the right inferior frontal gyrus (rIFG), two key nodes of the cortical inhibitory network thought to support the implementation of action stopping (Aron et al., 2007).

While NiBS intervention effects on *group* means are reported to be medium (Cohen’s *d* ≈ 0.4) (e.g., Hsu et al., 2015), correlations between *individual* neurophysiological and behavioural changes would be expected to be smaller due to measurement error and inter-individual variability. In the present context, a mechanistic link between preSMA-rIFG connectivity and response inhibition is theoretically plausible given the established causal role of this network (Aron et al., 2007; Schaum et al., 2021; Zhu et al., 2026). However, any underlying coupling is expected to be reduced at the observable level due to the imperfect reliability of both connectivity estimates and behavioural outcome measures (Hedge et al., 2018). To address this, we quantified change using RCIs, which incorporate measurement reliability when scaling pre- to post-change, providing a principled basis for testing whether neurophysiological and behavioural responses covary.

Through the application of meta-analytic techniques to pooled individual-level data, the available sample size (*N* = 69) provided statistical power of ∼0.47 for *r* = 0.2, ∼ 0.72 for *r* = 0.3, ∼0.93 for *r* = 0.4, and ∼0.99 for *r* = 0.5 (Figure 1) between within-subject changes in an electroencephalogram (EEG) connectivity measure (imaginary part of coherency, ImCoh) (Nolte et al., 2004) between preSMA and rIFG and corresponding behavioural outcomes (stop signal reaction time, SSRT). Given the expected medium to large effect at the group level (Cohen’s d ≈ 0.4; Hsu et al., 2015), and that individual-level correlations are typically attenuated relative group-level effects due to measurement (Hedge et al., 2018), any meaningful individual-level coupling would plausibly manifest as a moderate correlation (e.g., r ≈ 0.3–0.4). As such, with N = 69, the present sample was adequately powered to detect effects of this magnitude (power = 0.72–0.93), though power for a small effect (r = 0.2) remained modest (power = 0.47). While power curves of Pearson’s correlation are presented in Figure 1 for ease of interpretation, the primary analysis in the current paper used mixed-effects regression with participant included as a random factor. Since the effect of interest reflects the standardised association between ImCoh and SSRT change scores, power considerations are appropriately expressed in terms of detectable correlation magnitudes (Giner-Sorolla et al., 2024). This approach provides a principled test of whether individual tACS-induced changes in preSMA-rIFG connectivity are systematically related to individual changes in SSRT.

**Figure 1.**
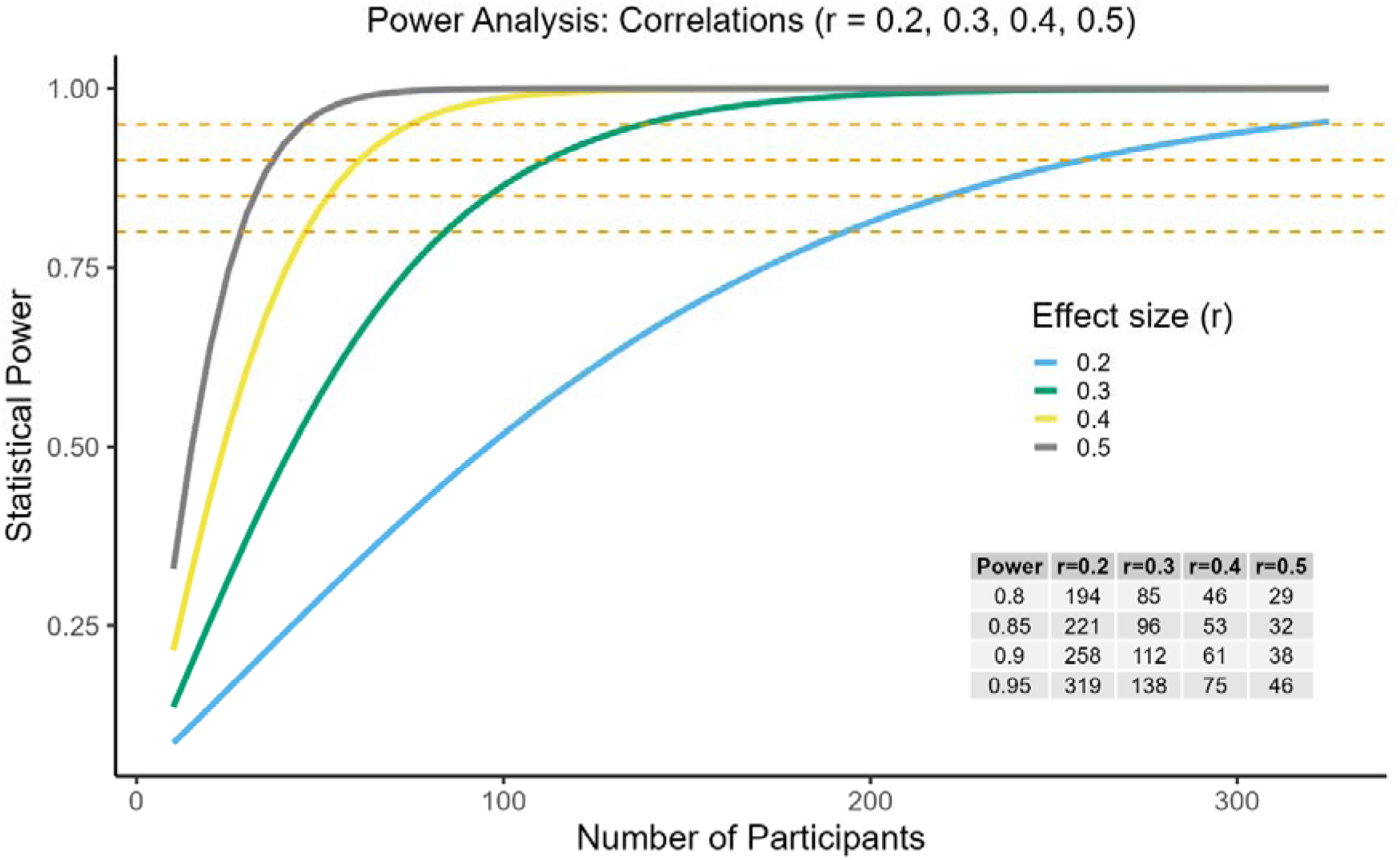
Power curves for detecting Pearson correlations of r = 0.2, 0.3, 0.4, and 0.5 at α = 0.05 (two-tailed). Coloured lines show statistical power across a range of sample sizes for each effect size. Horizontal red dashed lines indicate conventional target power levels (0.8, 0.85, 0.9, and 0.95). The embedded table in the bottom-right corner displays the exact number of participants required to achieve each target power for each effect size.

## Method

### Data Sources

Archived data were obtained from three independent NiBS experiments conducted in our research group between 2023 and 2025 (Fujiyama et al., 2023, 2025; Tan et al., 2025). Across these studies, data from a total of 69 young participants (38 males) were included, with a mean age of 24.3 years (SD = 5.4). All three studies involved 1) a pre-post design incorporating EEG ImCoh measure and SSRT, 2) tACS, and 3) individual-level baseline and post-intervention data suitable for multivariate analysis. All studies were approved by the Murdoch University Human Ethics Committee (2016/021 for Fujiyama et al., 2023 and Tan et al., 2025; 2022/186 for Fujiyama et al., 2025). Each study addressed a distinct research question, but all shared a common set of features comprising the use of young neurologically healthy participants, baseline assessment, tACS administration, and post-intervention assessment of both EEG and stop signal task (SST). Importantly, only active stimulation conditions/groups were included in the current analysis. A brief overview of each study is provided below and summarised in Table 1.

**Table 1:** Summary of studies with NiBS parameters, outcome measures, and relevant main outcomes

|  | Fujiyama et al. (2023) | Tan et al. (2025) | Fujiyama et al. (2025) |
| --- | --- | --- | --- |
| N (males) | 36 (22) | 18 (11) | 15 (5) |
| Mean Age (SD) | 23.7 (5.2) yrs | 23.6 (4.5) yrs | 24.5 (4.8) yrs |
| Design | Between-subject | Within-subject | Between-subject |
| NiBS parameters | Bifocal 1mA 20 Hz tACS targeting preSMA and rIFG for 15 min at 0° phase lag either during SST performance or prior to SST | Bifocal 1mA 20 Hz tACS targeting preSMA and rIFG for 15 min at 0° and 180° phase lag prior to SST | Bifocal 20 Hz tACS targeting preSMA (1mA) and rIFG (1.6mA) for 20 min at 0° phase lag during SST performance |
| Neurophysiological measure | Resting state ImCoh (128-channel HydroCel Geodesic Sensor Net, Magstim EGI, Eugene, OR) | Resting state ImCoh (128-channel HydroCel Geodesic Sensor Net, Magstim EGI, Eugene, OR) | Resting state ImCoh (128-channel HydroCel Geodesic Sensor Net, Magstim EGI, Eugene, OR) |
| Behavioural measure | SSRT: Two-choice RT task with stop signals to estimate SSRT, completed in two blocks at each assessment time point with 80 trials per block (60 Go, 20 Stop) | SSRT: A simple RT task with stop signals to estimate SSRT, completed in two blocks at each assessment time point with 100 trials per block (70 Go, 30 Stop) | SSRT): Two-choice RT task with stop signals to estimate SSRT, completed in two blocks at each assessment time point with 80 trials per block (60 Go, 20 Stop) |
| Main outcome | Offline stimulation improved SSRT despite no observable ImCoh changes. Online stimulation increased ImCoh without SSRT improvement | In-phase tACS increased ImCoh in both age groups, whereas anti-phase tACS reduced it only in older adults. Both conditions improved response inhibition in both age groups | No within-session change in ImCoh and SSRT |

Note: N in Fujiyama et al. (2023) and Fujiyama et al. (2025) excludes the participants assigned to the sham group, while the data from 15 older participants in Tan et al. (2025) were not included in the present study. From Tan et al. (2025), only data from 0° stimulation sessions were included for consistency across projects. For a multi-session study (Fujiyama et al., 2025), only Session 1 data were included in the pooled analysis. tACS = Transcranial alternating current stimulation, preSMA = pre-supplementary motor cortex, rIFG = right inferior frontal gyrus, M1= Primary motor cortex, SST = stop signal task; ImCoh = imaginary part of coherency; SSRT = stop signal reaction time

*Fujiyama et al. (2023):* This study investigated whether bifocal beta (20Hz) tACS (1mA) applied to the rIFG and preSMA was more effective during task performance (online) versus prior to task (offline) in modulating response inhibition. Fifty-three healthy adults (32 female, 18–35 years) received 15 minutes of phase-synchronised tACS or sham while performing a SST (i.e., between-subject design). Behavioural performance was measured using SSRT, and functional connectivity was measured using ImCoh. Offline tACS significantly improved response inhibition without overt changes in EEG connectivity, suggesting a post-stimulation plasticity effect. Online tACS increased post-stimulation functional connectivity but did not significantly improve inhibition. We included data from participants who underwent online or offline stimulation (n = 36), excluding those who received sham stimulation.

*Tan et al. (2025).* Bifocal beta (20 Hz) tACS over the rIFG and preSMA for 15 minutes was tested for its ability to modulate functional connectivity and response inhibition in older adults, alongside a young-adult comparison group, using a double-blind crossover design. Thirtyl7lthree healthy participants (15 older adults aged 61–79 years, 7 females; 18 young adults aged 18–34 years, 10 females) completed two stimulation sessions: inl7lphase tACS (0° phase offset) and antil7lphase tACS (180° offset). Note that we included only the young adult sample in the current paper.

Resting-state EEG assessed beta-band rIFG–preSMA connectivity, and inhibitory control was measured using SSRT. Inl7lphase tACS increased rIFG–preSMA connectivity in both age groups, whereas antil7lphase tACS reduced connectivity specifically in older adults. Despite these divergent neural effects, both stimulation conditions improved response inhibition in young and older adults. We included only data from 0° stimulation sessions in younger participants, excluding the 15 older participants.

*Fujiyama et al. (2025).* This study examined the effects of repeated bifocal beta tACS (20 Hz, 20 min) targeting the rIFG and preSMA over five sessions in 30 young adults (18–35 years, 10 femals) receiving either real or sham stimulation across 5 sessions over 2 weeks. Resting-state EEG assessed functional connectivity, response inhibition was measured using a SST, and simulated driving performance was evaluated before and after the intervention. Repeated tACS enhanced functional connectivity between preSMA and rIFG, but response inhibition and braking performance remained unchanged. General driving performance showed potential improvement, suggesting increased attentional capacity. For the present study, only session 1 data from the real stimulation group were included in the pooled analysis.

## Harmonisation of Measures Across Studies and Statistical Analyses

Data from three independent projects were combined to investigate brain-behaviour coupling following tACS. The RCI was used to quantify participant-specific change in preSMA–rIFG ImCoh and SSRT. For ImCoh, resting-state data from 3 min were segmented into 2-second epochs, yielding 90 repeated observations per time point (pre- and post-stimulation) per participant. For SSRT, 2 blocks per time point were collected, each containing left-stop and right-stop trials, yielding 4 repeated SSRT observations, yielding 2 SSRT estimates per block (left-stop, right-stop), per time point per participant. To account for the inherent measurement error of these metrics, the RCI was computed using the standard formula (i.e., *RCI* = *cx_post_* - *x_pre_*)/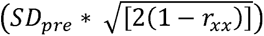, assuming a test-retest reliability *r_xx_ o*f .80. Positive RCI values indicate reliable improvement. Specifically, higher ImCoh reflects increased connectivity, while higher (sign-inverted) SSRT reflects faster stopping. SSRT scores were sign-inverted prior to RCI calculation so that lower raw values reflected positive change. To harmonise data across the three independent studies and control for study-specific variance, the resulting RCI scores were z-standardised within each study (mean = 0, SD = 1) prior to being pooled. These standardised RCI scores were then analysed using mixed-effects regression, with participant as a random intercept, to examine neurophysiological-behavioural associations. Although each participant contributed only one observation, a mixedl7leffects regression model with participant as a random intercept was used to formally account for potential nonl7lindependence of observations and to acknowledge participantl7lspecific variability. This approach provides a more conservative framework for estimating fixedl7leffects associations, even when the identifiability of the randoml7leffect variance is limited by the design.

Of note, study-wise z-standardisation controls for systematic between-study differences in RCI distribution (e.g., due to differences in sample size, participant characteristics, variations in study design), despite RCIs being already standardised for measurement reliability. In addition, to empirically evaluate the statistical limitations of ratio-based change scores, a sensitivity analysis was conducted, in which brain-behaviour associations were recomputed using post/pre ratio scores. As with the primary RCI-based analysis, ratio scores were z-standardised within each study prior to pooling, and associations were examined using the same mixed-effects regression analysis.

Statistical analyses and visualisations were conducted in R version 4.4.1 (R Development Core Team, 2021) within RStudio version 2025.09.2□+□418 (RStudio Team, 2020). A linear mixedl7leffects regression model was fitted using the lme4 package (Bates et al., 2015) with significance testing via lmerTest (Kuznetsova et al., 2017). Data manipulation and wrangling were performed using dplyr (Wickham et al., 2023) and tidyr (Wickham, Vaughan, et al., 2024), descriptive statistics were obtained using psych (Revelle, 2025), and graphical outputs were generated using ggplot2 (Wickham, 2016). Data import was handled with readr (Wickham, Hester, et al., 2024). Power analyses were conducted using the pwr package (Champely, 2020). All data and codes are publicly available on the Open Science Framework: https://osf.io/jfr3c.

## Results

### Summary of change scores

Across the full sample, the mean ImCoh RCI was 0.20 (SD = 2.03), and the mean SSRT RCI was 0.38 (SD = 1.28) (Table 2), indicating that, on average, participants showed modest improvements in ImCoh and SSRT (SSRT was sign-inverted prior to RCI calculation, hence positive scores reflect improvement). Study-specific means were small and variable: ImCoh RCIs ranged from −0.13 (Fujiyama et al., 2025) to 0.63 (Tan et al., 2025), and SSRT RCIs ranged from 0.12 (Fujiyama et al., 2023) to 0.79 (Tan et al., 2025), with Tan et al. (2025) showing the largest average improvements on both measures. These descriptive statistics highlight the substantial inter-individual variability, with some participants showing strong increases or decreases in connectivity and/or stopping performance, while others showed little or even opposite trends relative to their study average

**Table 2:** Overall and Study-Specific Reliable Change Indices (RCI) for ImCoh and SSRT

|  | n | Mean RCI | SD RCI | Min RCI | Max RCI |
| --- | --- | --- | --- | --- | --- |
| **ImCoh** |  |  |  |  |  |
| **overall** | 69 | 0.20 | 2.03 |  |  |
| Fuijyama et al. (2023) | 36 | 0.13 | 2.34 | -4.01 | 5.79 |
| Tan et al. (2023) | 18 | 0.63 | 2.06 | -3.51 | 3.97 |
| Fuijyama et al. (2025) | 15 | -0.13 | 0.94 | -1.57 | 1.99 |
| **SSRT** |  |  |  |  |  |
| **overall** | 69 | 0.38 | 1.28 |  |  |
| Fuijyama et al. (2023) | 36 | 0.12 | 1.41 | -2.13 | 4.43 |
| Tan et al. (2023) | 18 | 0.79 | 1.34 | -1.66 | 3.41 |
| Fuijyama et al. (2025) | 20 | 0.49 | 0.61 | -0.34 | 1.68 |

### Brain–behaviour association

Participant-level z-standardised RCI scores for preSMA–rIFG ImCoh and SSRT are shown in Figure 2. Scores were standardised within each study (mean = 0, SD = 1) to enable direct comparison across studies. Across the full sample (N = 69), the uncorrected correlation between ImCoh RCI and SSRT RCI was positive, but weak and nonl7lsignificant (r = 0.012, p = 0.92). A linear mixedl7leffects regression model (with participant ID as a random intercept and project as a fixed effect) indicated that ImCoh RCI was not a significant predictor of SSRT RCI (β = −0.05, SE = 0.10, t = −0.51, p = 0.99). There were no systematic differences among projects in SSRT RCI (all p > 0.9), and the fixedl7leffects component of the model accounted for only a negligible proportion of variance (marginal R² = 0.004).

**Figure 2.**
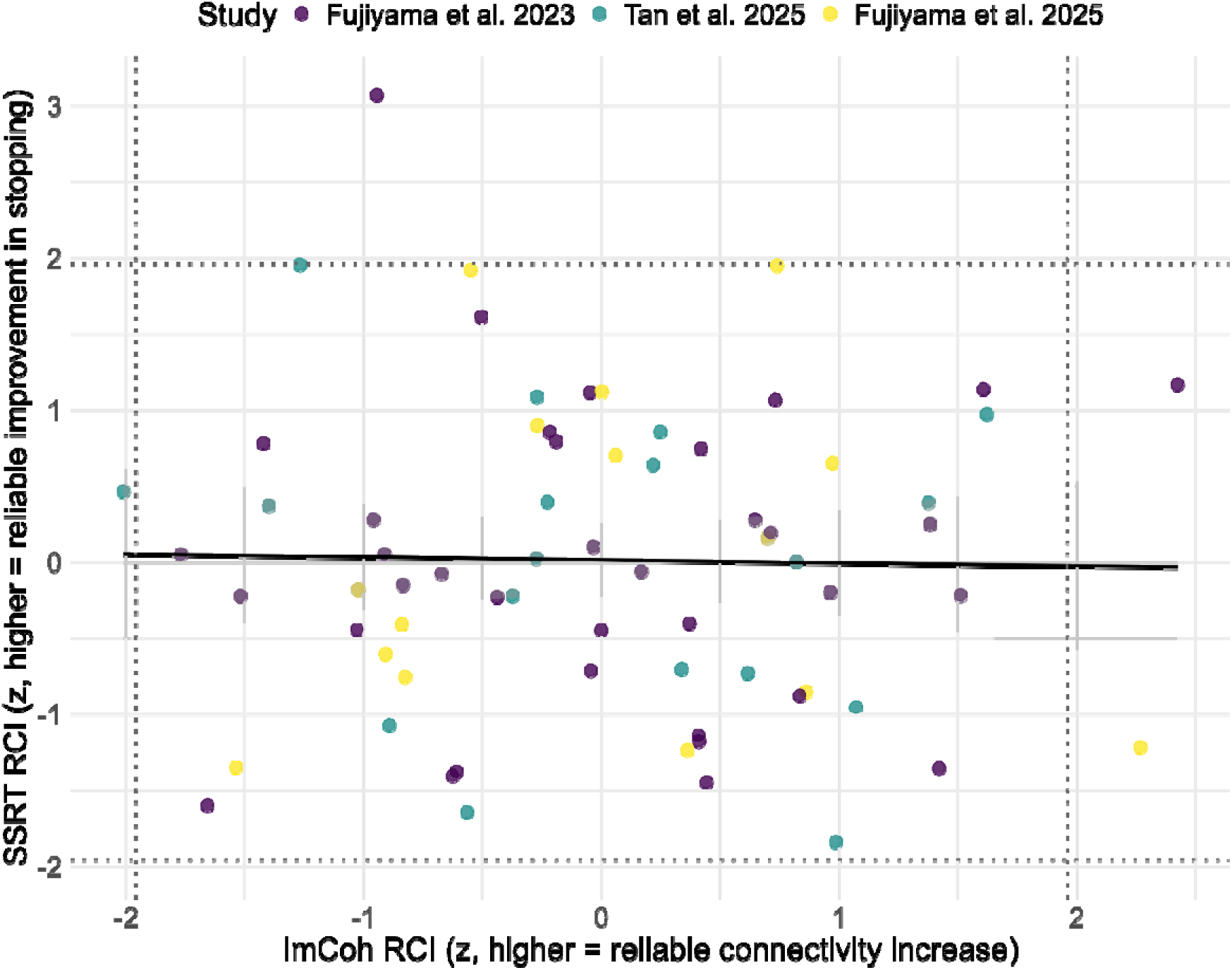
RCI-based brain–behaviour coupling across studies. Data points reflect participant-level z-standardised RCI scores for preSMA–rIFG ImCoh and SSRT (N = 69). RCI quantifies reliable intervention effects accounting for measurement error.

### Project-wise associations

Descriptive correlations between ImCoh and SSRT RCIs were small in magnitude and nonl7lsignificant across all three projects (Fujiyama et al., 2023: r = 0.04, p = 0.82; Tan et al., 2025: r = −0.20, p = 0.43; Fujiyama et al., 2025: r = 0.05, p = 0.86), with no evidence of consistent or statistically meaningful projectl7lspecific effects. Figure 2 illustrates substantial interl7lindividual variability in both neurophysiological and behavioural change, with the regression line reflecting only a weak overall directional covariation. Increases in ImCoh RCI were not robustly associated with greater improvements in SSRT RCI, despite the visual impression of modest positive trends in some individuals. Dotted lines indicate the ±1.96 RCI threshold for reliable change. Only seven participants exceeded this threshold on either ImCoh RCI or SSRT RCI, precluding meaningful subgroup analyses to test whether the association between ImCoh and SSRT differed for participants showing reliable improvement versus those showing little or no change.

Scores were standardised within projects (mean = 0, SD = 1) to enable cross-study comparability. Dotted lines indicate ± 1.96 RCI thresholds for reliable change. The regression line captures overall directional covariation across individuals (β = -.05, p = 0.99), indicating no meaningful relationship between tACS-induced changes in preSMA–rIFG connectivity and response inhibition at the individual level.

### Sensitivity analysis

To evaluate the robustness of the primary RCI-based findings and to empirically demonstrate the statistical limitations of ratio-based change scores described in the introduction, we recomputed brain–behaviour associations using post/pre ratio scores. The overall correlation between ImCoh and SSRT ratio scores was also negligible and non-significant (r = 0.014, p >0.9), mirroring the absence of a meaningful association observed with RCI-based scores (r = 0.012). This pattern is consistent with the expected noise amplification introduced by ratio-based measures, particularly when baseline variability is substantial. Projectl7lwise correlations using ratio scores were inconsistent in both direction and magnitude (Fujiyama et al., 2023: r = −0.08, p = 0.65; Tan et al., 2025: r = 0.18, p = 0.49; Fujiyama et al., 2025: r = 0.04, p = 0.88), similar to the variable pattern observed across projects using RCIl7lbased scores (Fujiyama et al., 2023: r = 0.04, p = 0.82; Tan et al., 2025: r = −0.20, p = 0.43; Fujiyama et al., 2025: r = 0.05, p = 0.86). A linear mixedl7leffects regression model using the same specification as the primary RCIl7lbased analysis indicated that the ImCoh ratio score was not a significant predictor of the SSRT ratio score (β ≈ 0, SE = 0.13, t = 0.11, p = 0.91), and the fixedl7leffects component of the model accounted for a negligible proportion of variance in SSRT change (marginal R² ≈ 0.03). As with the primary model, the solution was unstable and dominated by participantl7llevel variance, reinforcing the limitations of this approach in the current sample.

## Discussion

Using RCI in preSMA-rIFG functional connectivity (indexed with ImCoh) and response inhibition (indexed with SSRT) following 20 Hz tACS as an empirical example, this paper demonstrates a principled approach to analysing brain-behaviour coupling. This approach provides an alternative to the ratio-based change score commonly used in NiBS research, particularly in small and underpowered samples. To quantify individual change in a way that accounts for measurement reliability, change in both ImCoh and SSRT was operationalised using RCI. This approach provides a principled method for expressing pre- to post-change in standardised units while reducing the interpretational problems associated with ratio-based metrics (Kronmal, 1993). Mixed-effect regression modelling further accounted for repeated measures within participants and clustering across studies, addressing common limitations of prior post-hoc NiBS correlation analyses that are often underpowered for individual differences This paper demonstrates a principled approach to quantifying individual-level brain–behaviour coupling in NiBS research, using preSMA–rIFG functional connectivity (ImCoh) and response inhibition (SSRT) following 20 Hz tACS as an empirical testbed. Individual change was operationalised using RCI, which accounts for measurement unreliability and expresses pre-to-post change in standardised units, addressing a key limitation of the ratio-based change scores commonly used in the field (Kronmal, 1993). Mixed-effects regression modelling further accounted for clustering across studies while treating project as a fixed effect, enabling a more precise estimate of brain–behaviour coupling than any single contributing study could achieve alone (Button et al., 2013). Applying this framework to a pooled sample of N = 69 participants across three independent studies, however, revealed no meaningful association between tACS-induced changes in ImCoh and SSRT (r = 0.01, β = −0.05, p = 0.99, marginal R² = 0.004).

Project-wise correlations were small, non-significant, and inconsistent in direction, providing no support for a systematic individual-level brain–behaviour coupling under conventional 20 Hz tACS protocols. The sensitivity analysis using ratio-based change scores converged on the same null result (r = 0.014); while the two approaches yielded similar point estimates here, ratio scores showed severe distributional violations (Shapiro-Wilk W = 0.16 for SSRT ratio, p < .001) relative to the approximately normal RCI distributions (W = 0.975, p = .183), supporting the methodological rationale for preferring RCI on statistical grounds. Together, these results provide little support for the hypothesis that individuals showing greater tACS-induced increases in preSMA–rIFG beta synchrony also show proportionally greater improvements in stopping performance, and suggest that any true individual-level effect, if present, is likely to be small. Importantly, a null individual-difference association does not contradict group-level tACS effects reported in the contributing studies. These answer fundamentally different questions. Group-level analyses test whether stimulation shifts the average connectivity or behaviour across participants, whereas brain–behaviour coupling tests whether the magnitude of neurophysiological change predicts the magnitude of behavioural change at the individual level. The present results suggest these two levels of analysis can dissociate a distinction that warrants explicit consideration in future NiBS research.

Extensive imaging studies have established the preSMA and rIFG, together with the subthalamic nucleus, as critical hubs for response inhibition, with structural and functional connectivity during successful stops in SSTs (Aron, 2011; Jahfari et al., 2011). Functional communication between these regions is theorised to be essential for superior inhibitory control (Swann et al., 2012), with transcranial magnetic stimulation studies supporting their causal relationship (Chambers et al., 2006). In this context, the absence of a strong association is nonetheless interpretable within prevailing models, even though the magnitude of the effect was smaller than anticipated. Accordingly, the present results do not challenge the involvement of preSMA and rIFG in the response inhibition process, rather, they challenge the assumption that a simple linear connectivity metric provides a robust individual difference marker of behavioural gain.

While preSMA–rIFG beta synchronisation represents a plausible mechanistic target, ImCoh may not adequately capture the dynamic nonlinear coupling required for behavioural translation. This is consistent with imaging evidence showing context-dependent, sometimes paradoxical relationships between rIFG/preSMA activation and stopping performance, even at the group level stops (Tsvetanov et al., 2018; Wessel et al., 2012), challenging simplistic assumptions that higher connectivity necessarily produces better inhibition. The absence of directional consistency across projects (i.e., correlations ranging from r = −0.20 to r = 0.05), further suggests that any putative brain–behaviour association is not robust to study context and likely reflects sampling variability around a true effect close to zero.

Compounding this, even modest measurement unreliability in SSRT and resting-state connectivity would substantially attenuate observable associations in change score contexts. Together, these results underscore that neural measures alone provide limited traction for inferring individual behavioural change, and future work would benefit from designs that jointly assess both levels of analysis.

A key consideration in interpreting these findings is the substantial interindividual variability that characterises responses to NiBS. Converging evidence indicates that fewer than half of the participants show the expected neurophysiological response following NiBS, with marked variability also observed within individuals across sessions (Guerra et al., 2020; Vergallito et al., 2022), with some exceptions showing reasonable stability across sessions (e.g., Hinder et al., 2014). This variability is widely recognised as a major limitation for both the statistical sensitivity and clinical translation of NiBS. A recent review highlighted multiple interacting contributors, including stable factors such as anatomy, neurochemistry, and genetics, state-based influences, including hormonal fluctuations or substance intake, and contextual factors, including baseline capacity and task demands (Vergallito et al., 2022). Importantly, even when identical stimulation parameters are applied, participants can show different outcomes. Computational modelling has emerged as a tool to quantify how these identical parameters can produce varying electric field strengths and orientations at the target cortical site, providing insight into a source of interindividual variability (Evans et al., 2020). Such differences are linked to variability in both neurophysiological and behavioural outcomes across healthy and clinical populations (for review, see Hunold et al., 2023). However, these associations largely remain correlational, and greater stimulation intensity does not necessarily produce greater effects, challenging simple linear response assumptions (Lee et al., 2021).

Given this state-dependent variability, closed-loop NiBS approaches that continuously adapt stimulation to moment-to-moment brain state offer a principled alternative to fixed-parameter protocols (Frohlich & Townsend, 2021; Wansbrough, Tan, et al., 2024). Although empirical evidence remains limited (Stecher et al., 2021), such approaches are well-suited to addressing the inter- and intra-individual variability that constrains conventional protocols. The null association observed here may partly reflect the insensitivity of a static linear metric like ImCoh to the dynamic, state-dependent changes induced by tACS (Aydore et al., 2013; Bergmann, 2018), as well as the broader theoretical point that preSMA–rIFG communication is one component of an inhibitory network rather than a single causal determinant of behavioural improvement (Aron et al., 2014)

## Limitations and future perspectives

These present results indicate that changes in preSMA-rIFG functional interaction have weak predictive power of corresponding improvements in response inhibition. mCoh was selected as a conservative, volume-conduction-resistant index of phase synchronisation (Nolte et al., 2004), but the null result suggests no single static connectivity measure is likely to provide a robust individual-level marker of behavioural change following tACS. Future work should evaluate alternative neurophysiological markers and consider PCA-based change scores when trial-level data are available (Carson, 2024; Jolliffe & Cadima, 2016), which may improve stability when reliability is modest and reduce ratio artefacts. Additionally, the RCI computation assumed test-retest reliability of .80; future work should establish empirical reliability estimates for these specific measures.

Lastly, although outcome measures were harmonised via within-study standardisation, heterogeneity in stimulation parameters and task variants may have further attenuated any consistent brain-behaviour relationship. However, heterogeneity was relatively constrained: EEG hardware was identical across studies, and preprocessing pipelines were comparable, while tACS devices and electrode montages were aligned. Protocol differences in tACS were limited to modest variation in current intensity (1 mA in Fujiyama et al., 2023 and Tan et al., 2025; 1.6 mA for preSMA in Fujiyama et al., 2025) and stimulation duration (15 min vs 20 min), as well as task implementation (identical SST in Fujiyama et al., 2023 and 2025; simplified RT with stop signal in Tan et al., 2025). Critically, a linear mixed effects regression including study as a fixed effect still showed that ImCoh RCI did not significantly predict SSRT RCI (p > 0.9), suggesting that between-study heterogeneity played only a minor role in the null association.

## Conclusion

In summary, while prior work from these studies has reported group-level increases in preSMA–rIFG connectivity following 20 Hz tACS, the translation of these neurophysiological changes to individual-level improvement in response inhibition measured with SSRT remains weak and inconsistent. This modest brain-behaviour coupling, even in a pooled and harmonised dataset, underscores the substantial inter- and intra-individual variability that characterises NiBS responses. Beyond providing a cautionary note about interpreting post-hoc correlation in studies that are designed to capture pre- to post-changes, the present work offers a methodological benchmark: the combined use of RCI-based change scores and mixed-effects modelling enhances reliability and comparability across studies. At the same time, the results highlighted the need to identify more sensitive neural biomarkers, integrate multiple modal imaging, and explore state-dependent, i.e., closed-loop, stimulation protocols to improve behavioural outcomes and advance mechanistic understanding of neurophysiological responses to NiBS.

## Acknowledgments

This work was supported by the Neurotrauma Research Program (DoH20193370) awarded to H.F. and A.D.T. A.D.T. was supported by a Sarich Family Research Fellowship.

## References

1. Allison, D. B., Paultre, F., Goran, M. I., Poehlman, E. T., & Heymsfield, S. B. (1995). Statistical considerations regarding the use of ratios to adjust data. International Journal of Obesity and Related Metabolic Disorders.__: Journal of the International Association for the Study of Obesity, 19(9), 644–652.

2. Aron, A. R. (2011). From reactive to proactive and selective control: Developing a richer model for stopping inappropriate responses. Biological Psychiatry, 69(12), e55—68-e55—68. 10.1016/j.biopsych.2010.07.024

3. Aron, A. R., Durston, S., Eagle, D. M., Logan, G. D., Stinear, C. M., & Stuphorn, V. (2007). Converging evidence for a fronto-basal-ganglia network for inhibitory control of action and cognition. Journal of Neuroscience, 27(44), 11860–11864.

4. Aron, A. R., Robbins, T. W., & Poldrack, R. A. (2014). Inhibition and the right inferior frontal cortex: One decade on. Trends in Cognitive Sciences, 18(4), 177–185. 10.1016/j.tics.2013.12.003

5. Aydore, S., Pantazis, D., & Leahy, R. M. (2013). A note on the phase locking value and its properties. NeuroImage, 74, 231–244. 10.1016/j.neuroimage.2013.02.008

6. Bates, D., Mächler, M., Bolker, B., & Walker, S. (2015). Fitting Linear Mixed-Effects Models Using {lme4}. Journal of Statistical Software, 67(1), 1–48. 10.18637/jss.v067.i01

7. Benjamini, Y., Drai, D., Elmer, G., Kafkafi, N., & Golani, I. (2001). Controlling the false discovery rate in behavior genetics research. Behavioural Brain Research, 125(1–2), 279–284. 10.1016/s0166-4328(01)00297-2

8. Bergmann, T. O. (2018). Brain State-Dependent Brain Stimulation. Frontiers in Psychology, 9, 4–4. 10.3389/fpsyg.2018.02108

9. Bestmann, S., & Krakauer, J. W. (2015). The uses and interpretations of the motor-evoked potential for understanding behaviour. Experimental Brain Research, 233(3), 679–689. 10.1007/s00221-014-4183-7

10. Button, K. S., Ioannidis, J. P. A., Mokrysz, C., Nosek, B. A., Flint, J., Robinson, E. S. J., & Munafò, M. R. (2013). Power failure: Why small sample size undermines the reliability of neuroscience. Nature Reviews Neuroscience, 14(5), 365–376. 10.1038/nrn3475

11. Carson, R. G. (2024). A cogent technique to circumvent the use of (some) ratio measures in physiology. The Journal of Physiology, 602(19), 4713–4728. 10.1113/JP285214

12. Chambers, C. D., Bellgrove, M. A., Stokes, M. G., Henderson, T. R., Garavan, H., Robertson, I. H., Morris, A. P., & Mattingley, J. B. (2006). Executive “brake failure” following deactivation of human frontal lobe. Journal of Cognitive Neuroscience, 18(3), 444–455.

13. Champely, S. (2020). pwr: Basic Functions for Power Analysis. https://github.com/heliosdrm/pwr

14. Díaz-Francés, E., & Rubio, F. J. (2013). On the existence of a normal approximation to the distribution of the ratio of two independent normal random variables. Statistical Papers, 54(2), 309–323. 10.1007/s00362-012-0429-2

15. Evans, C., Bachmann, C., Lee, J. S. A., Gregoriou, E., Ward, N., & Bestmann, S. (2020). Dose-controlled tDCS reduces electric field intensity variability at a cortical target site. Brain Stimulation, 13(1), 125–136. 10.1016/j.brs.2019.10.004

16. Farahani, F., Kronberg, G., FallahRad, M., Oviedo, H. V., & Parra, L. C. (2021). Effects of direct current stimulation on synaptic plasticity in a single neuron. Brain Stimulation, 14(3), 588–597. 10.1016/j.brs.2021.03.001

17. Frohlich, F., & Townsend, L. (2021). Closed-Loop Transcranial Alternating Current Stimulation: Towards Personalized Non-invasive Brain Stimulation for the Treatment of Psychiatric Illnesses. Current Behavioral Neuroscience Reports, 8(2), 51–57. 10.1007/s40473-021-00227-8

18. Fujiyama, H., Bowden, V. K., Tang, A. D., Tan, J., Librizzi, E., & Loft, S. (2025). Repeated application of bifocal transcranial alternating current stimulation improves network connectivity but not response inhibition: A double-blind sham control study. *Cerebral Cortex (New York*, N.Y..__: 1991), *35*(5). 10.1093/cercor/bhaf110

19. Fujiyama, H., Hinder, M. R., Barzideh, A., Van de Vijver, C., Badache, A. C., Manrique-C, M. N., Reissig, P., Zhang, X., Levin, O., Summers, J. J., & Swinnen, S. P. (2017). Preconditioning tDCS facilitates subsequent tDCS effect on skill acquisition in older adults. Neurobiology of Aging, 51. 10.1016/j.neurobiolaging.2016.11.012

20. Fujiyama, H., Hyde, J., Hinder, M. R., Kim, S. J., McCormack, G. H., Vickers, J. C., & Summers, J. J. (2014). Delayed plastic responses to anodal tDCS in older adults. Frontiers in Aging Neuroscience, 6, 115–115. 10.3389/fnagi.2014.00115

21. Fujiyama, H., Williams, AlexandraG., Tan, J., Levin, O., & Hinder, M. R. (2023). Comparison of online and offline applications of dual-site transcranial alternating current stimulation (tACS) over the pre-supplementary motor area (preSMA) and right inferior frontal gyrus (rIFG) for improving response inhibition. Neuropsychologia, 108737–108737. 10.1016/j.neuropsychologia.2023.108737

22. Giner-Sorolla, R., Montoya, A. K., Reifman, A., Carpenter, T., Lewis, N. A. J., Aberson, C. L., Bostyn, D. H., Conrique, B. G., Ng, B. W., Schoemann, A. M., & Soderberg, C. (2024). Power to Detect What? Considerations for Planning and Evaluating Sample Size. Personality and Social Psychology Review.__: An Official Journal of the Society for Personality and Social Psychology, Inc, 28(3), 276–301. 10.1177/10888683241228328

23. Guerra, A., López-Alonso, V., Cheeran, B., & Suppa, A. (2020). Variability in non-invasive brain stimulation studies: Reasons and results. Non Invasive Brain Stimulation, 719, 133330. 10.1016/j.neulet.2017.12.058

24. Hedge, C., Powell, G., & Sumner, P. (2018). The reliability paradox: Why robust cognitive tasks do not produce reliable individual differences. Behavior Research Methods, 50(3), 1166–1186. 10.3758/s13428-017-0935-1

25. Hinder, M. R., Goss, E. L., Fujiyama, H., Canty, A. J., Garry, M. I., Rodger, J., & Summers, J. J. (2014). Inter- and intra-individual variability following intermittent theta burst stimulation: Implications for rehabilitation and recovery. Brain Stimulation, 7(3). 10.1016/j.brs.2014.01.004

26. Hsu, W.-Y., Ku, Y., Zanto, T. P., & Gazzaley, A. (2015). Effects of noninvasive brain stimulation on cognitive function in healthy aging and Alzheimer’s disease: A systematic review and meta-analysis. Neurobiology of Aging, 36(8), 2348–2359. 10.1016/j.neurobiolaging.2015.04.016

27. Hummel, F. C., Heise, K., Celnik, P., Floel, A., Gerloff, C., & Cohen, L. G. (2010). Facilitating skilled right hand motor function in older subjects by anodal polarization over the left primary motor cortex. Neurobiology of Aging, 31(12), 2160–2168. 10.1016/j.neurobiolaging.2008.12.008

28. Hunold, A., Haueisen, J., Nees, F., & Moliadze, V. (2023). Review of individualized current flow modeling studies for transcranial electrical stimulation. Journal of Neuroscience Research, 101(4), 405–423. Scopus. 10.1002/jnr.25154

29. Jacobson, N. S., & Truax, P. (1991). Clinical significance: A statistical approach to defining meaningful change in psychotherapy research. Journal of Consulting and Clinical Psychology, 59(1), 12–19. 10.1037//0022-006x.59.1.12

30. Jahfari, S., Waldorp, L., van den Wildenberg, W. P. M., Scholte, H. S., Ridderinkhof, K. R., & Forstmann, B. U. (2011). Effective Connectivity Reveals Important Roles for Both the Hyperdirect (Fronto-Subthalamic) and the Indirect (Fronto-Striatal-Pallidal) Fronto-Basal Ganglia Pathways during Response Inhibition. The Journal of Neuroscience, 31(18), 6891–6891. 10.1523/JNEUROSCI.5253-10.2011

31. Jasieński, M., & Bazzaz, F. A. (1999). The Fallacy of Ratios and the Testability of Models in Biology. Oikos, 84(2), 321–326. JSTOR. 10.2307/3546729

32. Jolliffe, I. T., & Cadima, J. (2016). Principal component analysis: A review and recent developments. Philosophical Transactions. Series A, Mathematical, Physical, and Engineering Sciences, 374(2065), 20150202. 10.1098/rsta.2015.0202

33. Kiernan, M., & Baiocchi, M. T. (2022). Casting New Light on Statistical Power: An Illuminating Analogy and Strategies to Avoid Underpowered Trials. American Journal of Epidemiology, 191(8), 1500–1507. 10.1093/aje/kwac019

34. Kronmal, R. A. (1993). Spurious Correlation and the Fallacy of the Ratio Standard Revisited. Journal of The Royal Statistical Society Series A-Statistics in Society, 156, 379–392.

35. Kuznetsova, A., Brockhoff, P. B., & Christensen, R. H. B. (2017). lmerTest Package: Tests in Linear Mixed Effects Models. Journal of Statistical Software, 82, 1–26. 10.18637/jss.v082.i13

36. Lebihan, B., Mobers, L., Daley, S., Battle, R., Leclercq, N., Misic, K., Wansbrough, K., Vallence, A.-M., Tang, A., Nitsche, M., & Fujiyama, H. (2025). Bifocal tACS over the primary sensorimotor cortices increases interhemispheric inhibition and improves bimanual dexterity. *Cerebral Cortex*, bhaf011–bhaf011. 10.1093/cercor/bhaf011

37. Lee, J.-H., Jeun, Y.-J., Park, H. Y., & Jung, Y.-J. (2021). Effect of Transcranial Direct Current Stimulation Combined with Rehabilitation on Arm and Hand Function in Stroke Patients: A Systematic Review and Meta-Analysis. *Healthcare (Basel*, Switzerland), 9(12). 10.3390/healthcare9121705

38. Nitsche, M. A., & Paulus, W. (2001). Sustained excitability elevations induced by transcranial DC motor cortex stimulation in humans. Neurology, 57, 1899–1901.

39. Nolte, G., Bai, O., Wheaton, L., Mari, Z., Vorbach, S., & Hallett, M. (2004). Identifying true brain interaction from EEG data using the imaginary part of coherency. Clinical Neurophysiology.__: Official Journal of the International Federation of Clinical Neurophysiology, 115(10), 2292–2307. 10.1016/j.clinph.2004.04.029

40. R Development Core Team. (2021). *R: A language and environment for statistical computing.* https://www.r-project.org/.

41. Reinhart, R. M. G., & Nguyen, J. A. (2019). Working memory revived in older adults by synchronizing rhythmic brain circuits. Nature Neuroscience, 22(5), 820–827. 10.1038/s41593-019-0371-x

42. Rektorová, I., Pupíková, M., Fleury, L., Brabenec, L., & Hummel, F. C. (2025). Non-invasive brain stimulation: Current and future applications in neurology. Nature Reviews Neurology, 21(12), 669–686. 10.1038/s41582-025-01137-z

43. Revelle, W. (2025). psych: Procedures for Psychological, Psychometric, and Personality Research. https://personality-project.org/r/psych/

44. RStudio Team. (2020). *RStudio: Integrated Development for R.* http://www.rstudio.com/.

45. Ryan, J. L., Eng, E., Fehlings, D. L., Wright, F. V., Levac, D. E., & Beal, D. S. (2023). Motor Evoked Potential Amplitude in Motor Behavior-based Transcranial Direct Current Stimulation Studies: A Systematic Review. Journal of Motor Behavior, 55(3), 313–329. 10.1080/00222895.2023.2184320

46. Schaum, M., Pinzuti, E., Sebastian, A., Lieb, K., Fries, P., Mobascher, A., Jung, P., Wibral, M., & Tüscher, O. (2021). Right inferior frontal gyrus implements motor inhibitory control via beta-band oscillations in humans. eLife, 10. 10.7554/eLife.61679

47. Stecher, H. I., Notbohm, A., Kasten, F. H., & Herrmann, C. S. (2021). A Comparison of Closed Loop vs. Fixed Frequency tACS on Modulating Brain Oscillations and Visual Detection. Frontiers in Human Neuroscience, *Volume 15-*2021. https://www.frontiersin.org/journals/human-neuroscience/articles/10.3389/fnhum.2021.661432

48. Swann, N. C., Cai, W., Conner, C. R., Pieters, T. A., Claffey, M. P., George, J. S., Aron, A. R., & Tandon, N. (2012). Roles for the pre-supplementary motor area and the right inferior frontal gyrus in stopping action: Electrophysiological responses and functional and structural connectivity. Neuroimage, 59(3), 2860–2870. 10.1016/j.neuroimage.2011.09.049

49. Tan, J., Iyer, K. K., Nitsche, M. A., Puri, R., Hinder, M. R., & Fujiyama, H. (2025). Dual-Site Beta tACS Over the rIFG and preSMA-Induced Phase-Specific Changes in Functional Connectivity but not Response Inhibition Performance in Older Adults. Psychophysiology, 62(5), e70060. 10.1111/psyp.70060

50. Tang, A. D., Bennett, W., Bindoff, A. D., Bolland, S., Collins, J., Langley, R. C., Garry, M. I., Summers, J. J., Hinder, M. R., Rodger, J., & Canty, A. J. (2021). Subthreshold repetitive transcranial magnetic stimulation drives structural synaptic plasticity in the young and aged motor cortex. Brain Stimulation, 14(6), 1498–1507. 10.1016/j.brs.2021.10.001

51. Tsvetanov, K. A., Ye, Z., Hughes, L., Samu, D., Treder, M. S., Wolpe, N., Tyler, L. K., & Rowe, J. B. (2018). Activity and connectivity differences underlying inhibitory control across the adult lifespan. The Journal of Neuroscience. 10.1523/jneurosci.2919-17.2018

52. Vergallito, A., Feroldi, S., Pisoni, A., & Romero Lauro, L. J. (2022). Inter-Individual Variability in tDCS Effects: A Narrative Review on the Contribution of Stable, Variable, and Contextual Factors. Brain Sciences, 12(5), 522. 10.3390/brainsci12050522

53. Vlachos, A., Müller-Dahlhaus, F., Rosskopp, J., Lenz, M., Ziemann, U., & Deller, T. (2012). Repetitive Magnetic Stimulation Induces Functional and Structural Plasticity of Excitatory Postsynapses in Mouse Organotypic Hippocampal Slice Cultures. The Journal of Neuroscience, 32(48), 17514. 10.1523/JNEUROSCI.0409-12.2012

54. Wansbrough, K., Marinovic, W., Fujiyama, H., & Vallence, A.-M. (2024). Beta tACS of varying intensities differentially affect resting-state and movement-related M1-M1 connectivity. Frontiers in Neuroscience, 18. https://www.frontiersin.org/journals/neuroscience/articles/10.3389/fnins.2024.142552 7

55. Wansbrough, K., Tan, J., Vallence, A.-M., & Fujiyama, H. (2024). Recent advancements in optimising transcranial electrical stimulation: Reducing response variability through individualised stimulation. Current Opinion in Behavioral Sciences, 56, 101360–101360. 10.1016/j.cobeha.2024.101360

56. Wessel, J. R., Danielmeier, C., Morton, J. B., & Ullsperger, M. (2012). Surprise and error: Common neuronal architecture for the processing of errors and novelty. The Journal of Neuroscience.__: The Official Journal of the Society for Neuroscience, 32(22), 528–7537. 10.1523/JNEUROSCI.6352-11.2012

57. Wickham, H. (2016). ggplot2: Elegant Graphics for Data Analysis. Springer-Verlag New York. https://ggplot2.tidyverse.org

58. Wickham, H., François, R., Henry, L., Müller, K., & Vaughan, D. (2023). *dplyr: A Grammar of Data Manipulation*. https://dplyr.tidyverse.org

59. Wickham, H., Hester, J., & Bryan, J. (2024). *readr: Read Rectangular Text Data*. https://readr.tidyverse.org

60. Wickham, H., Vaughan, D., & Girlich, M. (2024). *tidyr: Tidy Messy Data*. https://tidyr.tidyverse.org

61. Zehetmayer, S., Graf, A. C., & Posch, M. (2015). Sample size reassessment for a two-stage design controlling the false discovery rate. Statistical Applications in Genetics and Molecular Biology, 14(5), 429–442. 10.1515/sagmb-2014-0025

62. Zhu, T., Sack, A. T., & Leunissen, I. (2026). Phase-Specific Dual-Site Beta Transcranial Alternating Current Stimulation Differentially Influences Functional Connectivity Associated With Motor Inhibition Performance. Human Brain Mapping, 47(3), e70470. 10.1002/hbm.70470

63. Zimerman, M., Nitsch, M., Giraux, P., Gerloff, C., Cohen, L. G., & Hummel, F. C. (2013). Neuroenhancement of the aging brain: Restoring skill acquisition in old subjects. Annal of Neurology, 73(1), 10–15. 10.1002/ana.23761

